# TRAF6 modulates PD-L1 expression through YAP1-TFCP2 signaling in melanoma

**DOI:** 10.1101/2022.09.28.509909

**Authors:** Xiaoyan Liu, Linglu Wang, Yuhang Han, Hsiang-i Tsai, Fan Shu, Zhanxue Xu, Chao He, Haitao Zhu, Hongbo Chen, Fang Cheng

**Author notes:** **Correspondence to:** Haitao Zhu, Hongbo Chen, Fang Cheng E-mail addresses (Haitao Zhu), (Hongbo Chen), (Fang Cheng). Xiaoyan Liu and Linglu Wang contributed equally to this manuscript.

## Abstract

**Background:** Immunotherapy represented by the programmed death-1 (PD-1)/ligand 1 (PD-L1) monoclonal antibodies has led tumor treatment into a new era. However, the low overall response rate and high incidence of drug resistance largely damage the clinical benefits of existing immune checkpoint therapies. Recent studies correlate the response to PD-1/PD-L1 blockade with PD-L1 expression levels in tumor cells. Hence, identifying molecular targets and pathways controlling PD-L1 protein expression and stability in tumor cells is a major priority.

**Methods:** In this study, we first performed a Stress and Proteostasis CRISPR interference library-based screening to identify PD-L1 positive modulators. We then used *in vitro* and *in vivo* assays to investigate the biological function and mechanism of TRAF6 and its downstream YAP1/TFCP2 signaling in malignant melanoma.

**Results:** Here, we identified TRAF6 as a critical regulator of PD-L1 in melanoma cells. Suppression of TRAF6 expression down-regulates PD-L1 expression on the membrane surface of melanoma cells. We also found that PD-L1 protein abundance is regulated by YAP1/TFCP2 transcriptional complex. TRAF6 stabilizes YAP1 by K63 poly-ubiquitination modification, subsequently promoting the formation of YAP1/TFCP2 and PD-L1 transcription. Furthermore, inhibition of TRAF6 by Bortezomib enhanced cytolytic activity of CD8^+^ T cells by reduction of endogenous PD-L1. Notably, Bortezomib enhances anti-tumor immunity to an extent that is comparable to anti-PD-1 mAb therapies with no obvious toxicity.

**Conclusions:** These findings uncover a novel molecular mechanism for regulating PD-L1 protein abundance by a E3 ligase in cancer cells and reveal the potential of using TRAF6 inhibitors to stimulate internal anti-tumor immunological effect for TRAF6-PD-L1 overexpressing cancers.

## Background

Programmed death-ligand 1 (PD-L1) and its receptor programmed cell death protein 1 (PD-1) are the key targets to immune checkpoints blockade[1]. Monoclonal antibodies of PD-1/PD-L1 axis targeting have been approved for multi kinds of malignant carcinoma, which now serve as first-line clinical medication for malignant melanoma, pulmonary carcinoma, hepatoma, esophageal carcinoma, renal carcinoma and other advanced cancer and significantly prolong patient overall survival[2–4]. While prominent therapeutic effect has been obtained, low overall response rate of this immunotherapy gradually emerges as a notable obstacle, for only approximately 20%-40% patients can benefit from PD-1/PD-L1 immunosuppressive inhibitor treatment. Moreover, 15-35% benefited patients later obtain acquired resistance to immune checkpoint inhibitors, tumor relapse or progression after long-term PD-1/PD-L1 monoclonal antibodies treatment[5, 6].

Intrinsic and extrinsic factors such as tumor mutation burden, immune escape mechanisms, host status and immune infiltration are associated with the therapeutic effect of PD-1/PD-L1 monoclonal antibodies treatment[7], of which the expression level of PD-L1 on tumor cells is a key immune evasion indicator[4, 8, 9]. Various types of tumor cells highly express PD-L1 on their cell surfaces, which structurally binds to PD-1 located on the surface of T lymphocytes. This PD-1/PD-L1 specially binding impairs the activation and physiological function of cytotoxic T lymphocyte and assists tumor cells to evade from surveillance of immune system. In these kinds of tumor immune response reaction, inhibiting effect of PD-1/PD-L1 pathway is identified as a prominent rate-limiting step. Hence, immune checkpoint inhibitors targeting at interaction between PD-1 and PD-L1 potentiate hindrance of tumor evasion, reactivation of systemic anti-tumor reaction and rejuvenation of exhausted cytotoxic T lymphocytes in order to eradicate tumor cells. Yet, only part of PD-L1 positive patients benefit from PD-1/PD-L1 monoclonal antibodies treatment. Simultaneously, recent researches have uncovered the relationship between drug resistance of PD-1/PD-L1 monoclonal antibodies and PD-L1 expression of tumor. Multiple intrinsic factors including PDJ expansion (PD-L1/PD-L2/JAK2 gene amplification)[10], depletion of PTEN or mutation of PI3K/AKT[11], mutation of EGFR, overexpression of MYC, depletion of CDK5 [12] and 3’-UTR terminal truncations of PD-L1 contribute to continuous highly expression of PD-L1 on tumor cells, which subsequently put obstacles in the way of anti-tumor reaction and induce primary resistance to immune checkpoint inhibitors[13]. Recent studies indicate that not only cell membrane but also golgi apparatus, endosomes and microvesicles embodies PD-L1 molecular[14, 15]. These intracellular PD-L1 molecules promote tumor progression via re-transportation to cell surfaces[16] or secretion outwards in free form[17]. Meanwhile, a potential drug resistance mechanism of PD-1/PD-L1 inhibitors may result from the secretory PD-L1 molecules in peripheral blood which later reduce the therapeutic effect of immune checkpoint inhibitors by PD-L1 monoclonal antibodies consumption[17]. Another latent drug resistance mechanism of PD-1/PD-L1 immune checkpoint inhibitors is that PD-L1 could also be secreted via exosomes from tumor cells to tumor microenvironment and exert a negative influence on the anti-tumor reaction on cytotoxic CD8^+^ T lymphocytes[18]. So, it is of great importance to dig deeper in modulation mechanism of tumor PD-L1 molecules.

Human programmed death ligand (PD-L1), a transmembrane protein of 40kD, locates in chromosome 9p24.1, which is coded by CD274. Under the stimulation of interferon or other inflammatory factors, tumor cells rapidly raise PD-L1 expression and activate inhibitory PD-1/PD-L1 axis during tumorigenesis, which later elicits tumor immune escape. The abundance of PD-L1 in tumor microenvironment is modulated by a variety of factors, embracing alteration of genome (gene amplification and translocation)[10], modification of epigenetics (methylation of histone or CpG island, acetylation of histone and so on)[19, 20], transcriptional regulation (stimulation of inflammation and activation of PD-L1 transcriptional factors by carcinogenic signal)[21, 22], post-transcriptional regulation (miRNA, 3’-UTR and so on)[23] and post-translational modification (ubiquitination, phosphorylation, glycosylation, palmitoylation and so on)[24]. One group developed small molecule blockers and competitive peptides that inhibit palmitoylation of PD-L1 based on the mechanism of palmitoylation up-regulation of PD-L1 expression to clear endogenous PD-L1 of tumors[25]. Other studies found that metformin can down-regulate endogenous PD-L1 of tumors, and showed synergistic antitumor activity after combined treatment with CTLA-4 antibody[26]. These reports suggest that the discovery of new regulatory molecules of tumor PD-L1 provides new ideas for tumor immunotherapy.

In this study, we performed a Stress and Proteostasis CRISPR interference (CRISPRi) library-based screening to identify a new PD-L1 positive modulator, TRAF6. Suppression of TRAF6 expression down-regulates PD-L1 expression on the membrane surface of melanoma cells. Further mechanistic study indicates that TRAF6 stabilizes YAP1 by K63 poly-ubiquitination modification and promotes the formation of a transcriptional complex, YAP1/TFCP2 in the nucleus. And this stabilization increases the transcriptional expression of PD-L1 and subsequently promotes tumor immune escape. In addition, TRAF6 inhibition reduced PD-L1 levels in tumor cells and effectively enhanced cytolytic activity of CD8^+^ T cells. Importantly, TRAF6 inhibitor Bortezomib reduced numbers of tumor-infiltrating lymphocytes, enhances tumor regression and dramatically improves overall survival rates in murine metastatic melanoma models. Taken together, this study identifies a previously unrecognized regulatory pathway of PD-L1 expression in tumor cells and highlights a promising therapeutic target to enhance anti-tumor immunity.

## Materials and Methods

### Cell lines and cell culture

Cells used in this study (A2058, SK-MEL-5, HEK293T, B16F10) were obtained from the American Type Culture Collection (ATCC). Cells were cultured in high glucose Dulbecco’s Modified Eagle Medium (DMEM, GIBCO) containing 1% penicillin/streptomycin (P/S, Corning) and 10% fetal bovine serum (FBS, Sigma) at 37°C in a 5% CO_2_ atmosphere.

### Western blotting

Cells were completely lysed with ultrasonic for three times and five seconds at every turn using strong RIPA buffer (150mM NaCl, 1.0% NP-40, 0.5% sodium deoxycholate, 0.1% SDS, 50mM pH=8.0 Tris-HCl) with Roche complete protease inhibitor. Supernatants obtained from 1min centrifugation at 12,000 r were quantified using Bio-Rad Protein Assay Dye Reagent and successively heated to 100°C in 1×loading buffer for 5 min. Then denatured protein samples were separated by SDS-PAGE and transferred to PVDF membrane. Membranes were blocked by 5% non-fat milk in 1×TBST (1×TBS+0.1% Tweens) for 1h and incubated with primary antibodies at 4°C overnight. Reactive bands on membranes were visualized via ECL chemiluminescence kit following incubation with secondary antibodies conjugated to horseradish peroxidase for 2h.

### CRISPR-Cas9 based high-throughput screen

The mixture of dCas9 plasmids and packaging plasmids were firstly transfected into HEK 293T cells, and then supernatants containing the virus were collected 36 h after transfection. Melanoma cells were cultured with the lentiviral solution for 24 h in the presence of 1 μg/mL polybrene (Abmole, USA). After Blasticidin screening for 72 h, melanoma cell lines A2058-dCas9-KRAB stably expressing dCas9-KRAB were constructed. A2058-dCas9-KRAB cell lines were infected with Stress and Proteostasis-human subpooled sgRNA library (Cat#83973, Addgene), with MOI=0.3 and selected with puromycin (2 μg/mL) for 72 h, then the cells infected by library were collected 9 days after infection. The collected cells were divided into two parts averagely, one for control, one for enrichment of PD-L1low melanoma cells. Genomic DNA of enriched PD-L1low cells and unsorted cells were extracted respectively to amplify barcoded sgRNA-containing fragments of genomic DNA by two rounds of PCR. PCR products were first separated with 1.5% agarose gel, and then purified DNA fragments were subsequently recovered to perform high-throughput sequencing (illumina).

### Flow cytometry

Cells were washed with PBS and stained for special antibody in Cell Staining Buffer on ice for 15∼20 min. Then pellets uniformly resuspended in PBS were further analyzed on a MoFlo XDP flow cytometer.

### RNA isolation and qPCR analysis

About 2×10^6^ cells or abraded spleen tissues were lysed with 1ml RNAiso Plus. Cracked samples were added with 0.2ml trichloromethane and centrifuged at 12,000r for 10min at 4°C to obtain three distinct layers. The top layer was gained and added with isopropanol of same volume. After centrifugation, RNA precipitates were washed in 75% ethanol twice and dried in fume hood for 5∼10min. Purified RNA were further redissolved in DNAase/RNAase-free water. Then 1μg of entire RNA were reversely transcripted to cDNA using EasyScript® All-in-One First-Strand cDNA Synthesis SuperMix kit. Real-time PCR were executed on Roche Light Cycler 96 quantitative PCR instrument with SYBR Green. Primers were listed in Supplementary Table 2.

### Co-immunoprecipitation assay (Co-IP)

All samples were treated with 15μM MG132 for 6h before collecting to retain integrity of the ubiquitin chain. After removing culture medium and washing with PBS for 2 times, cells were lysed in RIPA buffer for 30 min. Total amount of 1mg cellular protein were then incubated with primary antibodies at 4°C overnight. A/G-Sepharose beads were added to capture antigen-antibody complex at 4°C for subsequent analysis by Western blotting.

### Chromatin immunoprecipitation (ChIP)

Cells were fixed with 1% formaldehyde at RT for 15 min. Glycine was added to quench cross-linking reactions. After washing with PBS, the cells were collected and resuspended in ChIP lysis buffer. The sheared chromatin was incubated with the indicated antibodies or control IgG and the protein A/G beads (Invitrogen, USA) were then added to capture protein-DNA complexes. The protein-DNA complexes were released from the beads by using 5M NaCl at 65°C for at least 6 h. Finally, the DNA was purified by phenol/chloroform/isoamyl alcohol (25:24:1, v/v/v) extraction and ethanol precipitation. The DNA was collected to perform a qPCR analysis.

### TMT proteomics

For TMT labeling, cell samples were firstly lysed with ultrasonic to extract proteins. Then, proteins were reduced by 5 mM dithiothreitol (DTT), alkylated by 11 mM iodoacetamide (IAA), and lysed by trypsin. After that, proteins were desalted with a Strata xc18 SPE column and vacuum-dried to obtain the peptides, and the peptides were labeled with TMT isobaric tags according to the instructions of the TMT kit.

For HPLC fractionation, the peptides were separated into 60 fractions by a gradient of 8%-32% acetonitrile (pH 9.0) for 60 min. Then, the peptides were vacuum freeze-dried after merging into 14 fractions. The peptides were isolated by an Agilent 300 Extend C18 column (5 μm diameter, 4.6 mm inner diameter, 250 mm length).

The peptides were separated by using an EASY-nLC 1000 UPLC after dissolving in solvent A. Then, the peptides were ionized by using NSI source and mass spectrometry was performed on a Q Exactive™ Plus. LC-MS/MS experiments were performed on the Orbitrap Elite LC-MS/MS.

### JASPAR based analysis

Possible binding sequences of TFCP2 before the transcriptional start sites of CD274 were predicted by JASPAR and sorted by Relative Score. Search TFCP2 in the JASPAR database (https://jaspar.genereg.net/), then input a FASTA-formatted sequence before the transcriptional start sites (TSS) of *CD274* which range from 0bp to 6000bp to scan with selected matrix models. Relative profile score threshold was set as 80%.

### GEO database analysis

The “PD-L1 totalnorm.txt” file of GSE129968 in the GEO database (htp://www ncbi nm.nh.gov/geo/) was downloaded and then the data of the unsorted cells (control group) and the PD-L1low cells (experimental group) were selected to merge the sgRNAs scores corresponding to each gene. Subsequently, fold change was calculated by dividing the experimental group by the control group and standardized in the form of log2. After that, P value was calculated and standardized in the form of -log10(P-Value). Graphic analyses were performed using Microsoft Excel 2010. The normalized file “series matrix file” from the melanoma microarray data GSE15605 was downloaded and average was performed when a gene corresponds to multiple gene probes. The Pearson correlation coefficient between genes and PD-L1 expression in 12 metastatic melanoma tissues were then calculated. Graphic analyses were performed using GraphPad Prism 8. Input the genes with a correlation coefficient greater than 0.4 and the expression matrix (GSE15605) of PD-L1 in 16 normal skin tissues and 12 metastatic melanoma tissues into the TB tool software to draw a heat map, then set the Row Scale to normalized and perform normalization to compare gene expression trends.

### Luciferase reporter assay

Different promoters or enhancers of PD-L1 were constructed into the vector, then the mixture of target plasmid, PD-L1 reporter gene plasmid and Renilla luciferase pRL-TK plasmid (4:1:1) were transfected into HEK 293T cells. After 48 h, the cells were harvested in the lysis buffer and the fluorescence value was measured by a dual luciferase assay kit (Promega, Madison, WI, USA).

### PBMCs and T cells isolation

Peripheral blood was contributed by healthy donors. Peripheral blood mononuclear cells (PBMCs) were isolated by density gradient centrifugation. Firstly, the blood sample was carefully layered onto Ficoll Paque Plus (GE Healthcare, USA) and centrifuged at 350 × g for 1h at 22°C to attain four layers. The cloudy-looking layer containing the PBMCs was collected and transferred to a new centrifuge tube. Then the PBMCs were incubated with red blood cell lysis buffer (Solarbio, China) and were washed with RPMI 1640 for three times. T cells were separated from PBMCs using a MojoSort human CD3^+^ T cell isolation kit (Biolegend, USA).

### *In vitro* co-culture system

Human CD3^+^ T cells isolated from peripheral blood were activated by 10μg/ml previously plate-bound anti-CD3 and 2μg/ml anti-IL-2for 48h. Melanoma cells were seeded into 24-well flat bottom plates one day in advance Then activated human CD3^+^ T cells were added to the 24-well plates with drugs treatment. After 12h, T cells were collected to detect the level of cytokines by flow cytometry and melanoma cells were collected to detect the level of apoptosis (Annexin V-APC/PI) by flow cytometry.

### CFSE staining

Blood-isolated CD3^+^ T cells were incubated with 5μM CFSE at 37°C for 20 min and washed with PBS for five times. Subsequently, samples were activated with 10μg/ml previously plate-bound anti-CD3 and 2μg/ml anti-IL-2 for 72h. After activation, CD3^+^T cells were treated with drugs and collected at different times points to detect cell proliferation by flow cytometry.

### Melanoma lung metastasis model

Laboratory mice were purchased from Experimental Animal Center of Sun Yat-sen University (animal certification number was 2001A032). Animal experiments were approved by the Animal Ethics Committee of Sun Yat-sen University, China. The approval number was No. 2018000577. To perform melanoma lung metastasis,2×10^6^ B16F10 mouse melanoma cells were injected into tail vein of C57BL/6 mice. A total of twenty mice were randomly separated into four well-distributed groups: group 1(normal control without cell injection, n=5), group 2(PBS, n=5), group 3(0.2mg/kg Bortezomib, n=5), group 4(100ug/kg anti-PD-1, n=5). Operated mice were injected with Bortezomib in tail vain or anti PD-1 in abdomina every three days for five times following mice melanoma cell injection. Half of mice were sacrificed and collected samples on D18, then the lungs were photographed, weighed, and the melanoma nodules in the lungs were calculated. While the other were recorded body weight every day and survival situations until death. The dissected lung tumor nodules were stained with hematoxylin-eosin (H&E) for histological analysis. The lungs of different groups of mice were cut into paraffin-embedded sections or frozen sections, and the expression level of TRAF6, PD-L1 and other proteins in the dissected lungs were detected by immunohistochemistry and western blot. In addition, the lungs, spleen and peripheral lymph nodes were sufficiently grinded and filtered through 70 μm filters to obtain single cell clusters for detecting CD4^+^/CD8^+^ and the level of IFNγ and GranzymeB by flow cytometry. Mouse serum were collected for detecting the level of IL-2 and IFN γ by ELISA.

### Score of tumor nodes

Tumor nodes were measured by vernier scale. The grade of tumor nodes was defined as below. I tumor nodes, diameter<0.5mm; II tumor nodes, 0.5mm≤diameter<1.0mm; III tumor nodes, 1.0mm≤diameter<2.0mm; IV tumor nodes, diameter≥0.5mm. The scores were counted by I tumor nodes*1+II tumor nodes*2+III tumor nodes*3+IV tumor nodes*4.

### Histology analysis

At the end of the experiment, the mouse recipients were sacrificed, and tissue samples were collected and submerged with 4% paraformaldehyde in 15 mL centrifugal tubes for fixation. The samples were sectioned to 4 µm thickness and stained with H&E or specific antibodies for immunochemistry staining following standard procedures. Histopathological state was observed by fluorescence microscopy with 20x magnification.

### Statistical analysis

Three independent sample replicates were carried out for each experiment unless stated otherwise. All data are expressed as the mean ± SD. Data analysis and processing were performed using GraphPad Prism Ver 7.0 software (GraphPad Software). The statistical significance between the two groups was measured using one-way analysis of variance (ANOVA) followed by Tukey’s test was performed, and statistical significance was indicated (*, *p* ≤ 0.05; **, *p* ≤ 0.01; ***, *p* ≤ 0.001).

## Results

### Identification of TRAF6 as a PD-L1 regulator by Proteostasis CRISPRi screening

To identify positive regulators of constitutive or induced cell surface PD-L1 expression, we performed a large-scale genetic CRISPR interference (CRISPRi) screen. A pooled human CRISPRi sgRNA lentivirus library with 16615 single-guide RNAs (sgRNAs) targeting 3323 human proteostasis-related genes together with dCas9-KRAB enzyme were introduced into in A2058 melanoma cells by lentiviral transduction. Deep sequencing of the sgRNAs integrated into genomic DNA from control cells, PD-L1^low^cells and PD-L1^high^ cells was subsequently performed (Figure 1A). We also reanalyzed a similar genome-Scale CRISPR Knock-out screening of PD-L1 regulator in lung cancer and comparison of the sequencing data led to the identification of sgRNAs that were enriched in PD-L1^low^cells, and the gene targets of the enriched sgRNAs by Volcano Plot analysis are potential PD-L1 negative regulators (Figure 1B). Intriguingly, among the 231 screen hits, 4 genes (TRAF6, PSMD8, CPNE9, SENP3) were identified and validated in our CRISPRi screening and genome-scale CRISPR Knock-out screening of PD-L1 regulator in lung cancer (Figure 1C). As all the above-mentioned genes are based on the data obtained from cancer cell lines, we then analyzed the correlation between these genes and PD-L1 expression in patient melanoma samples. Of the 231 genes obtained in Figure 1B, 144 were unregulated in metastatic melanoma tissues, including PD-L1 (Figure 1D). We further analyzed the expression matrix of top 26 genes with highest R values in the Pearson correlation analysis together with PD-L1 in 16 normal skin tissues and 12 metastatic melanoma tissues. Interestingly, TRAF6 also came out of genes with highest expression level in metastatic melanoma when compared to normal skin tissues (Figure 1E). As a summary, TRAF6 emerged as a gene essential for cell surface PD-L1 expression upon proteostasis CRISPRi high-throughput screening combining data from GEO databases of melanoma and lung cancer. To validate the role of TRAF6 in melanoma PD-L1 expression, we knocked down TRAF6 via short-hairpin RNA (shRNA) in A2058 and SK-MEL-5 melanoma cell lines. Consistent with the screening results, knocking down TRAF6 reduced PD-L1 expression, both in total protein level confirmed by western blot and cell surface level confirmed by flow cytometry (Figure 1F-G). Exogenous expression of TRAF6 in A2058 and SK-MEL-5 cells unregulated cell surface and total PD-L1 protein expression (Figure 1H-I), further confirming the positive role of TRAF6 modulating PD-L1 protein expression. TRAF6 belongs to E3 ubiquitin ligase family and conveys polyubiquitin chains to substrate protein. Besides Lys48-connected polyubiquitination and proteosome-dependent degradation, TRAF6 also catalyzes Lys63 (K63) -connected ubiquitination of substrates and enhances substrates stability, which further affects protein localization and multiple signaling pathways[27] In order to determine whether TRAF6 stabilizes PD-L1 directly by post-transcriptional K63 ubiquitinated modification, we expressed exogenous TRAF6 in 293T cells and found no obvious signal of PD-L1 K63 ubiquitination by immunoprecipitation (IP) (Figure 1J). Therefore, we supposed that TRAF6 modulate PD-L1 at transcriptional level. To test this hypothesis, we performed real-time quantitative PCR in TRAF6 knocked-down and TRAF6 over-expressing melanoma cell lines. Analysis of relative PD-L1 mRNA levels indicated that silencing TRAF6 reduced PD-L1 mRNA expression, whereas TRAF6 over-expression reduced PD-L1 mRNA expression (Figure 1K-L), indicating TRAF6 regulated PD-L1 expression in melanoma cells via transcriptional modulation.

**Figure 1.**
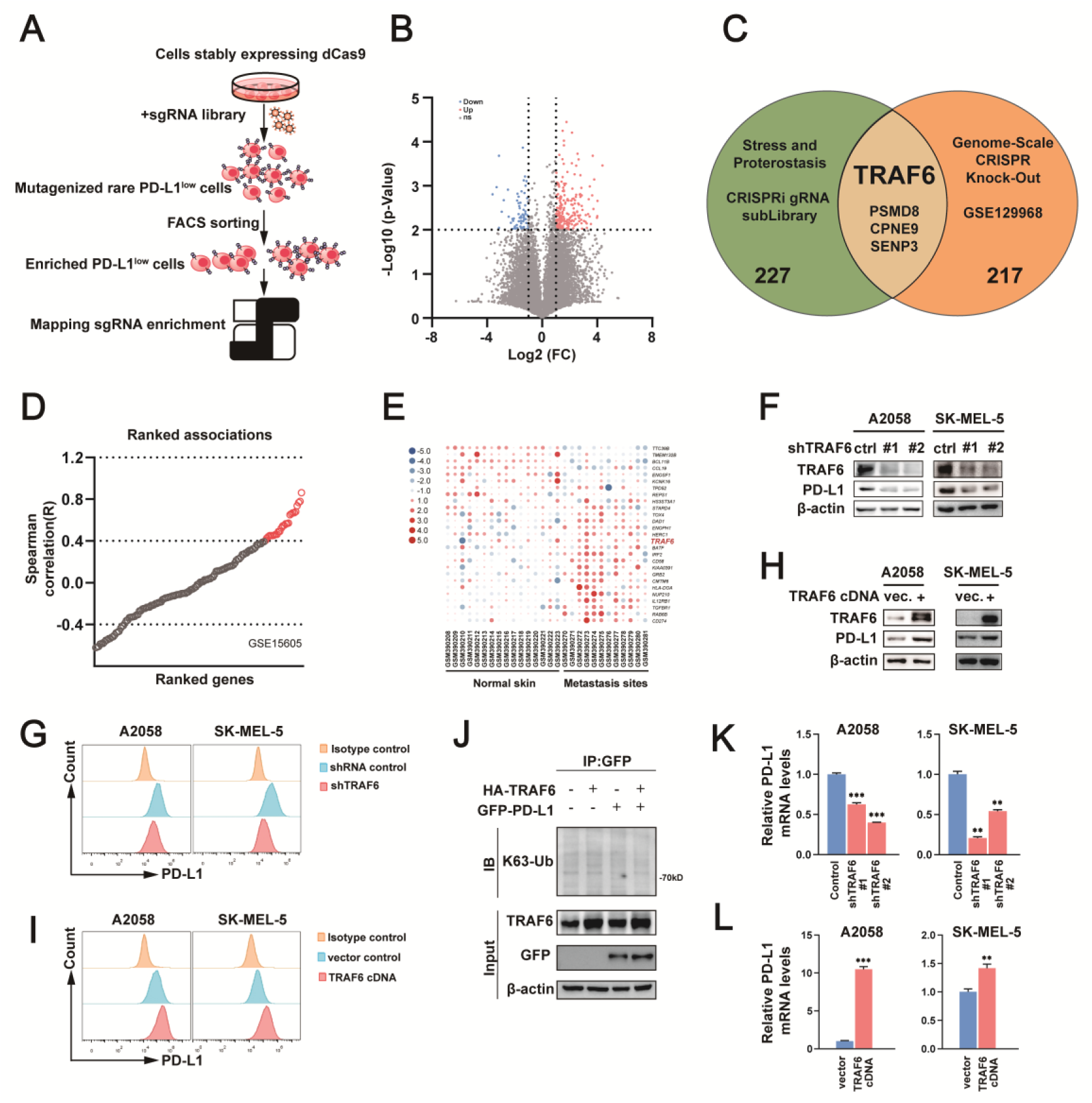
TRAF6 serves as a novel PD-L1 positive modulator in melanoma cells. **(A)** Schema of CRISPRi library-based screening. **(B)** Volcano Plot analysis of 20667 genes in GSE129968. Genes with the -Log10 (P-values) greater than or equal to 2 and with the Log2 (FC) greater than or equal to 1 were targeted as potential positive modulator. **(C)** Venn diagram presents the overlap of 221 targeted genes from GSE129968 and 231 positive-selected genes from Stress and Proteostasis CRISPRi gRNA subLibrary screening. Data of Stress and Proteostasis CRISPRi gRNA subLibrary screening were analyzed by MAGeCK. **(D)** The Pearson correlations (R) between 143 genes and PD-L1 expression in twelve cases of metastatic melanoma were calculated (GSE15605). The genes were arranged from small to large in order of R. R with greater than 0.4 represented medium and strong correlation. **(E)** Heat map of 26 genes and *CD274 (PD-L1)* expression in sixteen normal skin tissues and twelve metastatic melanoma tissues (GSE15605). After in-line standardization, red and blue respectively represented high and low expression. **(F-G)** PD-L1 expression in the TRAF6-knockdown A2058 and SK-MEL-5 cells analyzed by (F) western blot and (G) flow cytometry. **(H-I)** PD-L1 expression in the TRAF6-overexpressing A2058 and SK-MEL-5 cells analyzed by **(H)** western blot and **(I)** flow cytometry. **(J)** Immunoblots of anti-GFP immunoprecipitation in the HEK 293T cells co-expressing with HA-TRAF6 and GFP-PD-L1. **(K)** Transcriptional levels of PD-L1 in the TRAF6-knockdown A2058 and SK-MEL-5 cells. Mean with s.e.m.; n=3; **p<0.01; ***p<0.001; analyzed by Brown-Forsythe and Welch ANOVA tests. **(L)** Transcriptional Levels of PD-L1 in the TRAF6-overexpressing A2058 and SK-MEL-5 cells. Mean with s.e.m.; n=3; **p<0.01; ***p<0.001; analyzed by Unpaired t tests.

### TRAF6 regulates PD-L1 via TFCP2 binding to PD-L1 Promoter

To dig out the underlying mechanism of TRAF6 in PD-L1 transcriptional modulation, we first conducted tandem mass tag (TMT)-based quantitative proteomics analysis of TRAF6-depleted (shTRAF6) and control cell samples to identify TRAF6-regulating transcriptional factors (Figure 2A). We took the intersection of 6839 differentially expressed proteins identified in the proteomics analysis and 1665 transcriptional factors (TFs) documented in the Animal Transcription Factor DataBase (AnimalTFDB) to obtain 265 overlapping proteins (Figure 2B), 37 of which were obviously downregulated in TRAF6-depleted cells (Figure 2C). To further determine which factors may be more possibly involved in PD-L1 regulation, we utilized two human metastatic melanoma cohorts (*UCLA, Cell 2016; DFCI, Science 2015*) [28, 29] from cBioportal database to calculate correlation of mRNA expression of above 37 TFs coefficient with PD-L1 expression (Figure 2D-E). Sorted by relevance, TFCP2 was considered as the top candidate connecting TRAF6 and PD-L1 in these analyses. In line with above data analysis, we confirmed that knocking down TRAF6 significantly reduced TFCP2 expression (Figure 2F). Next, we explored whether TFCP2 is the transcriptional factor of PD-L1. We first confirmed that TFCP2 upregulated PD-L1 mRNA and consequent protein expression in TFCP2-overexpressed melanoma cells (Figure 2G), implying that TFCP2 could act as the direct transcriptional factor of PD-L1. Then JASPAR, a website containing transcriptional factor binding sites, was utilized to forecast DNA motifs on PD-L1 promoter interacted with TFCP2 (Figure 2H). Top ten predicted sequences scored by JASPAR were aligned to bases before PD-L1 transcriptional start site (TSS) (Figure 2I, Supplementary Table 3). Subsequently, TFCP2 binding affinity of these 10 sequences were verified by ChIP assay using anti-TFCP2 antibody, and Pro1 (AAACTGGATT) emerged as the most promising binding region with the highest enrichment (Figure 2J). RT-PCR further confirmed the relative enrichment of predicted sequences by TFCP2-CHIP conducted in A2058 cell line (Figure 2K). Located at -72bp before PD-L1 TSS, TFCP Pro1 possibly serves as a promoter for PD-L1 transcription. To test this possibility, we conducted dual luciferase reporter assay and revealed that overexpressing TFCP2 stimulates PD-L1 transcriptional activity when co-transfected with PD-L1 TFCP Pro1-luc plasmid (Figure 2L), confirming that TFCP2 served as a transcription factor and bound to PD-L1 promoter for transcription enhancement.

**Figure 2.**
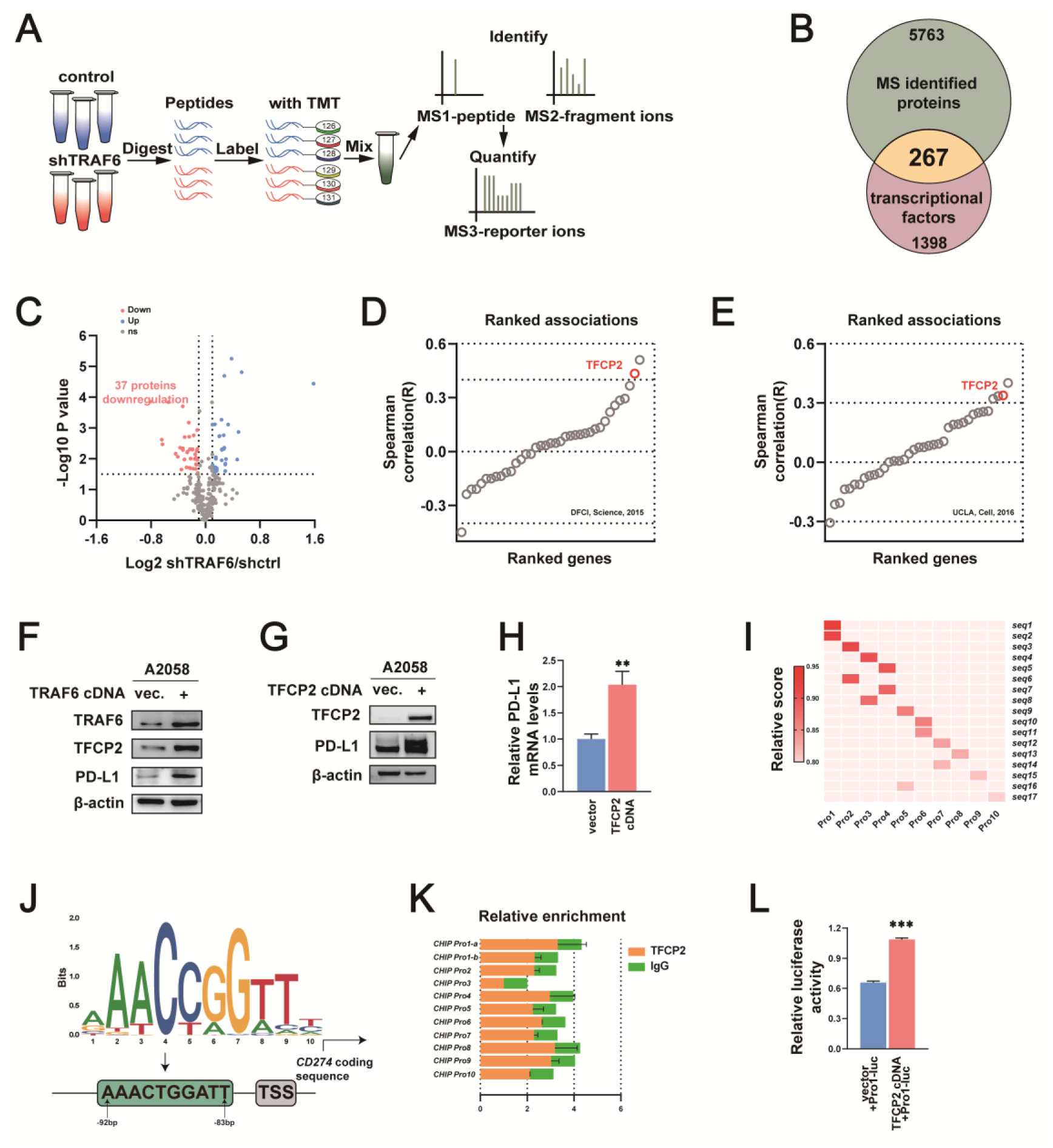
TRAF6 modulates PD-L1 transcriptional levels via TFCP2 binding to PD-L1 Promotor. **(A)** Schema of Tandem Mass Tag (TMT)-based quantitative proteomics. Analysis was conducted by comparing the TRAF6-kncokdown (shTRAF6) and the control (shctrl) A2058 cells. **(B)** Venn diagram presents the overlap of results from (A) and the transcriptional factors library. 6030 proteins were identified by TMT-based quantitative proteomics. The transcriptional factors library embodying 1665 proteins was documented in the Animal Transcription Factor DataBase. 267 transcriptional factors were overlapped. **(C)** Volcano Plot analysis of 267 overlapped proteins. 37 downregulated proteins were defined as -Log10 (P-values) greater than 1.5 and with the Log2 (FC) less than or equal to -0.1. **(D-E)** Ranked associations of 37 downregulated proteins with PD-L1 mRNA expression from cBioportal database. Plot (D) was analyzed with *DFCI, Sciences, 2015* and plot (E) was analyzed with *UCLA, Cell, 2015*. **(F)** Immunoblots analyzed the TRAF6-overexpressing A2058 cells. **(G)** Immunoblots analyzed the TFCP2-overexpressing A2058 cells. **(H)** Transcriptional levels of PD-L1 in the TFCP2-overexpressing A2058 cells. Mean with s.e.m.; n=3; **p<0.01; analyzed by Unpaired t tests. **(I)** Possible binding sequences of TFCP2 before the transcriptional start sites (TSS) of *CD274* predicted by JASPAR and were sorted by Relative Score. Sequences before TSS of *CD274* ranged from 0bp to -6000bp. Relative profile score threshold was set as 80%. **(J)** The best-matched DNA binding motif of TFCP2 predicted by JASPAR and could be aligned to -83bp to -92bp before TSS of *CD274*. **(K)** RT-PCR showed the relative enrichment of predicted sequences by TFCP2-CHIP conducted in A2058. Mean with s.d.; n=2. **(L)** Relative luciferase activity of HEK 293T samples after indicated transfection. Mean with s.e.m.; n=6; ***p<0.001; analyzed by Unpaired t tests.

### TRAF6 stimulates YAP1 K63 ubiquitination to modulate PD-L1

In order to understand how TRAF6 located in the cytosol compartment regulates TFCP2 transcriptional function in the nucleus, we first would rule out possible direct regulatory mechanism between TRAF6 and TFCP2. As expected, we confirmed that TRAF6 overexpression had no effect on TFCP2 K63 ubiquitination, indicating there may be an indirect regulatory mechanism between TRAF6 and TFCP2 (Figure 3A). Therefore, we performed the protein-protein interaction enrichment analysis using the BioGrid database. YAP1, a transcriptional co-activator that mainly interacts with TEAD family transcription factors to promote gene expression of the Hippo pathway, was identified as a TFCP2 co-factor via a WW-PSY motif to stimulate the transcription of YAP1 downstream proto-oncogenes (Figure 3B) [30]. Interestingly, YAP1 is a reported transcriptional co-factor of PD-L1 in many types of cancer[31–34]. Thus, we hypothesized that YAP1 serve as a transcriptional co-factor in TRAP6-TFCP2-mediated PD-L1 regulation. We first confirmed YAP1-TFCP2 interaction via co-immunoprecipitation assay in an over-expressing cell system (Figure 3C). To further confirm whether YAP1 plays a critical role in regulating TRAF6 mediated PD-L1 protein abundance, we applied genetic methods to deplete endogenous YAP1 in cells. The results showed that depletion of endogenous YAP1 using sgRNAs resulted in a dramatic downregulation of PD-L1 protein levels (Figure 3D). Importantly, ectopic expression of TRAP6 elevated the PD-L1 protein level in sgControl cells, but much milder in sgYAP1 cells (Figure 3D), suggesting TRAP6-mediated upregulation of PD-L1 is largely dependent on the YAP1 genetic status. In keeping with the results that YAP1 mediates TRAF6-PD-L1 pathway, ectopic expression of TRAP6 increased both YAP1 and PD-L1 protein abundance (Figure 3E). Furthermore, depletion of YAP1 in TRAP6-overexpressing cells dramatically reduced PD-L1 expression (Figure 3E), supporting that YAP1 support an essential intermediate role in TRAF6-PD-L1 modulation. In line with these results, inhibition of TRAF6 by shRNA lentivirus downregulated the protein levels of YAP1 and PD-L1, whereas ectopic expression of TRAF6 enhanced both YAP1 and PD-L1 expression in both A2058 and SK-MEL-5 melanoma cell lines (Figure 3F-G), suggesting that TRAF6 is a positive regulator of YAP1 and PD-L1. Since TRAF6 has the potential to stabilize substrate protein stability by K63 polyubiquitination, we then investigated whether TRAF6 could influence YAP1 protein stability utilizing cycloheximide (CHX) chase experiment. As expected, results from CHX assay also showed TRAF6 has a positive effect on YAP1 protein stability (Figure 3H-I), indicating YAP1 may be substrate for TRAF6 K63 ubiquitination. It is known that TRAF6 contains a N-terminal RING finger, five Zinc-finger repeats, a coiled-coil structure (trimer formation domain), as well as TRAF-C domain which can directly bind substrates (Figure 3J). Importantly, TRAF6 functions as an E3 ubiquitin ligase catalyzing K63-linked ubiquitination of target proteins, which modulate protein activity, localization, and its interaction with other proteins[35]. As the C70 residue of TRAF6 ring finger domain is critical for its ubiquitin-ligase activity, we inactivated its RING domain (TRAF6^C70A^) using point mutation to prevent its E3 ligase function. TRAF6 overexpression (TRAF6^WT^) stimulated K63 but not K48-linked ubiquitination, which is often associated with proteasome degradation of endogenous YAP1. Furthermore, TRAF6^C70A^ completely reversed the effect of TRAF6^WT^ on YAP1 K63 ubiquitination, demonstrating that the inhibition of TRAF6 ubiquitin-ligase activity attenuated the K63 ubiquitination of YAP1 as well as YAP1 protein stability (Figure 3K). Intriguingly, a very recent study has demonstrated that IL-1 induces YAP1 nuclear localization and protein stability by TRAF6-mediated K63-linked poly-ubiquitination of YAP1 at K252 in macrophages [36]. In this study, K321 and K497 site of YAP1 reported AS critical sites of its K63-linked ubiquitination by mass spectrometry analysis. We then generated two YAP1 mutants with the lysine 321 and 497 residue replaced by arginine (YAP1^K321R^ and YAP1^K497R^) and examined the TRAF6-mediated K63 poly-ubiquitination level of these two mutants in HEK 293T cells co-transfected with HA-TRAF6 vector (Figure 3J). YAP1^WT^ and YAP1^K321R,^ but not YAP1^K497R^ was poly-ubiquitinated in cells co-transfected with TRAF6 (Figure 3M). These data suggest that TRAF6-mediated K63-linked ubiquitination was located at K497 of YAP1.

**Figure 3.**
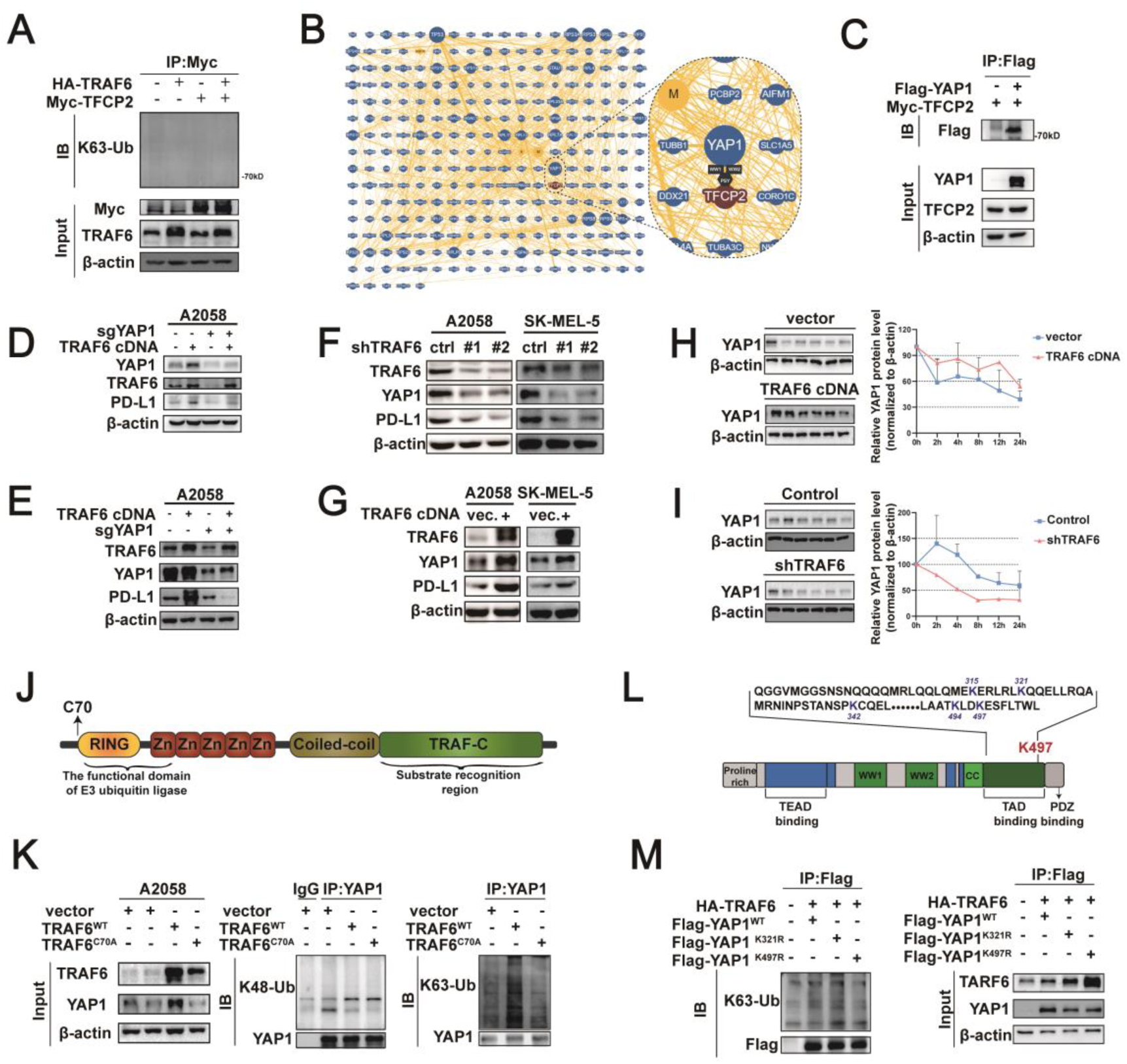
TRAF6 stimulates YAP1 K63 ubiquitination to modulate PD-L1. **(A)** Immunoblots of anti-Myc-Tag immunoprecipitation in the 293T cells co-expressing with HA-TRAF6 and Myc-TFCP2. **(B)** Interaction network of TFCP2 analyzed by BioGRID. Possible protein as interactors were presented as blue nodes. Minimum evidence was set as 3 and the network only show physical interaction and high throughput interaction. **(C)** Immunoblots of anti-Flag-Tag immunoprecipitation in the 293T cells co-expressing with Flag-YAP1 and Myc-TFCP2. **(D-E)** Immunoblots analyzed A2058 cells (D) knocked YAP1 off and then transfected with TRAF6 cDNA and (E) transfected with TRAF6 cDNA and then knocked YAP1 off. **(F-G)** Immunoblots analyzed A2058 and SK-MEL-5 cells with (F) TRAF6 knockdown and (G) TRAF6 overexpression. **(H-I)** Immunoblots analyzed A2058 cells after CHX treatment with (H) TRAF6 knockdown and (I) TRAF6 overexpression. **(J)** Basic structure of TRAF6 and the C70 site on the RING region is crucial for the activity of the E3 ubiquitin ligase. **(K)** Immunoblots of anti-YAP1 immunoprecipitation in the A2058 cells after indicated transfection. **(L)** Possible ubiquitinated lysine residues on the transactivation domain of YAP1. **(M)** Immunoblots of anti-Flag-Tag immunoprecipitation in the HEK 293T cells with indicated transfection.

### Inhibition of TRAF6 activated CD8^+^ T cells to eliminate melanoma *in vitro* and *in vivo*

To decipher the pathological significance of increased TRAF6 level in melanoma, we first examined if silencing TRAF6 in melanoma cells is sufficient to promote CD8^+^ T cell cytotoxicity. To do so, we developed a co-culture system in which we first stimulated CD3^+^ T cells, sorted from human peripheral blood mononuclear cells with anti-CD3 and IL-2 to induce cells to proliferate and produce cytotoxic cytokines. These stimulated T cells were then co-incubated with A2058 melanoma cells for 48h before *in vitro* cytotoxicity of both T cells and melanoma cells was detected via flow cytometric analysis (Figure 4A). The results of Annexin^+^/PI^+^ double staining showed that silencing TRAF6 accelerated melanoma cells entering into apoptosis (Figure 4B), which is consistent with earlier reports of TRAF6 required for cancer cell proliferation[37–39]. Silencing TRAF6 further induced apoptotic melanoma cells in the presence of stimulated CD8^+^ T cells (Figure 4B), indicating that inhibition of TRAF6 activates CD8^+^ T cells to eliminate melanoma cells. To achieve more evidence of TRAF6 as a potential target in tumor immunotherapy, we first optimized the concentration of the TRAF6 inhibitor Bortezomib by western blotting and determined 20 nM was optimal to achieve best inhibition of TRAF6 protein expression (Figure 4C). As expected, YAP1 and PD-L1 were also downregulated in melanoma cells when treated with 10nM-30nM Bortezomib (Figure 4C-D), supporting Bortezomib mediated TRAF6-YAP1-PD-L1 signaling in cancer cells. To test if Bortezomib mediated endogenous PD-L1 degradation is sufficient to promote CD8^+^ T cell cytotoxicity, we compared CD8^+^ T cell activation and IFN-gamma secretion when pretreated with Bortezomib or with PD-1 antibody in the co-culture system. Intriguing, although IFN-γ^+^ CD8^+^ T cell number was less in Bortezomib group (Figure 4E), in the T cell/tumor cell co-culture system, tumor cell killing indicated by Annexin V/PI double-staining was higher in Bortezomib group when compared in PD-1 antibody group (Figure 4F), suggesting that Bortezomib has a more dramatic effect in enhancing T cell killing of tumor cells than PD-1 antibody. Notably Bortezomib has no effect in T cell proliferation measured by flow cytometry analysis of CFSE (Figure 4G). Our results above demonstrated that inhibition of TRAF6 by genetic depletion or the pharmacological inhibitor, Bortezomib, upregulates PD-L1 protein levels and consequently enhances T cell induced apoptosis of tumor cells. Based on the molecular mechanism study, we hypothesized that inhibition of TRAF6 might enhances T cell-mediated anti-tumor immunity *in vivo*. To test this hypothesis, we utilized the syngeneic mouse B16F10 pulmonary metastasis melanoma model to examine if Bortezomib affected tumor growth and mice survival to a comparable level to anti-PD-1 mAb (Figure 4H). Strikingly, our results showed that TRAF6 inhibitor Bortezomib and anti-PD-1 mAb groups showed prominently weight gain and prolonged overall survival compared with PBS group (Figure 4I-J). In addition, Bortezomib significantly suppressed tumor development, evidenced by the reduced scores of melanoma nodules as well as smaller tumor size and areas in dissected lung tissues (Figure 4L-K). Hematoxylin&eosin and immunohistochemistry analysis also revealed increased density of Ki67+ proliferating cells and infiltrated T cells in both Bortezomib group and anti-PD-1 mAb group at a similar level (Figure 4M). These results support that the TRAF6 inhibitor treatment might reprogram an inflamed tumor microenvironment evidenced by upregulation of PD-L1, which enhances the tumor infiltrating CD8^+^ cytotoxic T cells to enable the PD-1/PD-L1 blockade *in vivo*.

**Figure 4.**
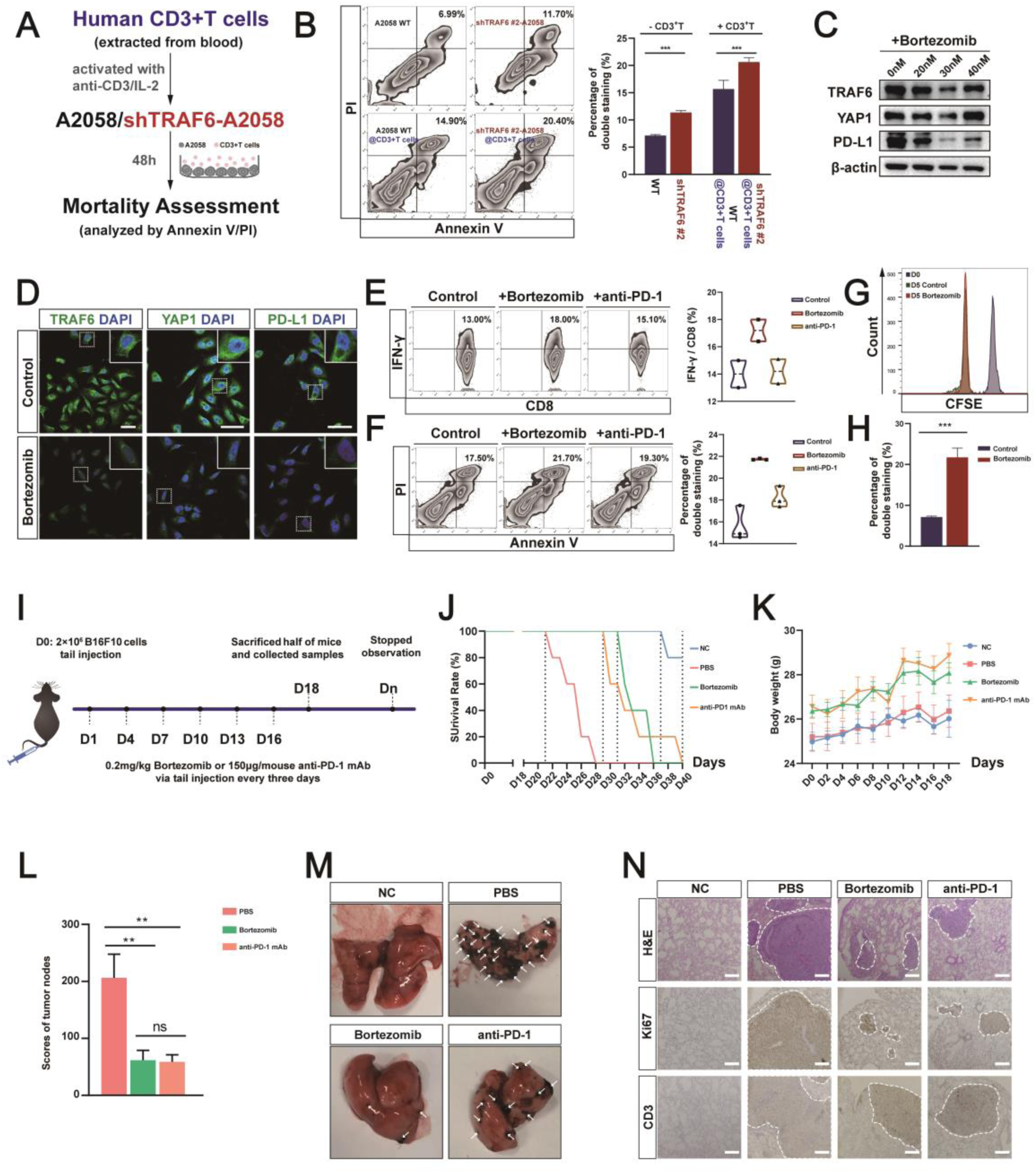
Inhibition of TRAF6 restrains tumor growth and elevates CD8^+^ T cells immune function. **(A)** Schema of the co-culture system consisting of A2058 cells and human CD3^+^T cells. A2058 cells were wild type (WT) or TRAF6 knockdown (shTRAF6). **(B)** Representative flow cytometric images and quantification of A2058 cells in the co-culture systems after Annexin V-APC /PI staining. Mean with s.e.m.; n=3; ***p<0.001; analyzed by Two-way ANOVA tests. **(C)** Immunoblots analyzed the A2058 cells treated with Bortezomib. **(D)** Representative immunofluorescent images of A2058 cells treated with 30nM Bortezomib. **(E-F)** Representative flow cytometric images and quantification of cells in the co-culture system after indicated treatment. Bortezomib was 30nM and anti-PD-1 mAb was 0.5μg/ml. (E) IFN-γ^+^ CD8^+^ cytotoxic T cells. Mean with s.d.; n=2. (F) A2058 cells stained by Annexin V-APC/PI. Mean with s.e.m.; n=3. **(G)** CFSE staining of T cells treated with 30nM Bortezomib analyzed by flow cytometry. **(H)** Quantification of A2058 cells stained by Annexin V-APC/PI after 30nM Bortezomib treatment for 12h. Mean with s.e.m.; n=3; ***p<0.001; analyzed by Unpaired t tests. **(I)** Regimens in the pulmonary metastasis melanoma model on the C57/BL6 mice. NC, no tumor-bearing mice; PBS, administrated with PBS; Bortezomib, administrated with 0.2mg/kg Bortezomib; anti-PD-1, administrated with 150μL/mouse anti-PD-1 mAb. **(J-K)** Life index of mice with indicated treatments. (J) Survival rates, n=6. (K) Body weight, mean with s.e.m.; n=5. **(L-M)** Tumor progression of mice with indicated treatments. (L)Score of tumors nodes, mean with s.e.m.; n=5; **p<0.01; ns, no significant; analyzed by One-way ANOVA tests. (M) Representative images of lung. **(N)** H&E staining and immunochemistry staining of the lung after indicated treatments.

We further investigated whether down-regulation of TRAF6 could affect the stability of YAP1, so as to decrease PD-L1 on tumor cell surface and elevate T cell toxicity targeting melanoma cells. Significant down-regulation of TRAF6, YAP1 and PD-L1 were observed in tumor tissues treated with the Bortezomib compared with control treatment (NC), as determined by immunostaining and western blot (Figure 5A-B). Intriguing, anti-PD-1 mAb treatment had no effect on TRAF6 expression, but moderately downregulated YAP1 and PD-L1 expression in the tumor region, which can be explained by preference of eliminating tumor cells with active YAP1-PD-L1 signaling by anti-PD-1 mAb treatment. Analysis of infiltrated immune cells demonstrated that Bortezomib treatment could significantly increase the percentage of CD8^+^ T cells, but not the CD4+ cells in tumor-infiltrating lymphocytes (Figure 5C). To further address whether the TRAF6 inhibitor affects the activation of tumor-infiltrating T cells, we also detected the T-cell activation maker, Granzyme B (GzmB) and IFN-γ, in infiltrated CD8^+^ T cells in syngeneic mouse melanoma model. Our results showed that both Bortezomib and anti-PD-1 mAb treatment significantly elevated the expression of GzmB and IFN-γ on infiltrated CD8^+^ T cells (Figure 5D-E). Additionally, we further dissected cervical lymph node, spleens, and peripheral lymph node to explore Bortezomib-induced potential negative effects on normal immunity. Although CD8^+^ T lymphocytes of Bortezomib group slightly augmented in these tissues, no apparent difference was observed (Figure 5F). Together, our results demonstrate that high expression of TRAF6 in tumors without treatment might enhance TRAF6-mediated K63-linked ubiquitination and stabilization of YAP1. YAP1 ubiquitination consequently promotes TFCP2 activity to transcribe its direct target gene PD-L1, leading to the suppression of tumor specific T-cell activity *in vitro* and *in vivo* (Fig. 5G, left panel). Inhibition of TRAF6 using the inhibitor Bortezomib can can reverse this process to reduce endogenous PD-L1, thus activating tumor-infiltrating killer CD8^+^ T cells and thus cause obvious cytolytic effect against melanoma cells (Fig. 5G, right panel). Notably, the therapy with TRAF6 inhibitor Bortezomib has similar efficacy to anti-PD-1 mAb treatment in metastatic melanoma model, indicating that potential superiority of TRAF6 as novel target to release constraints of immune checkpoint PD-L1.

**Figure 5.**
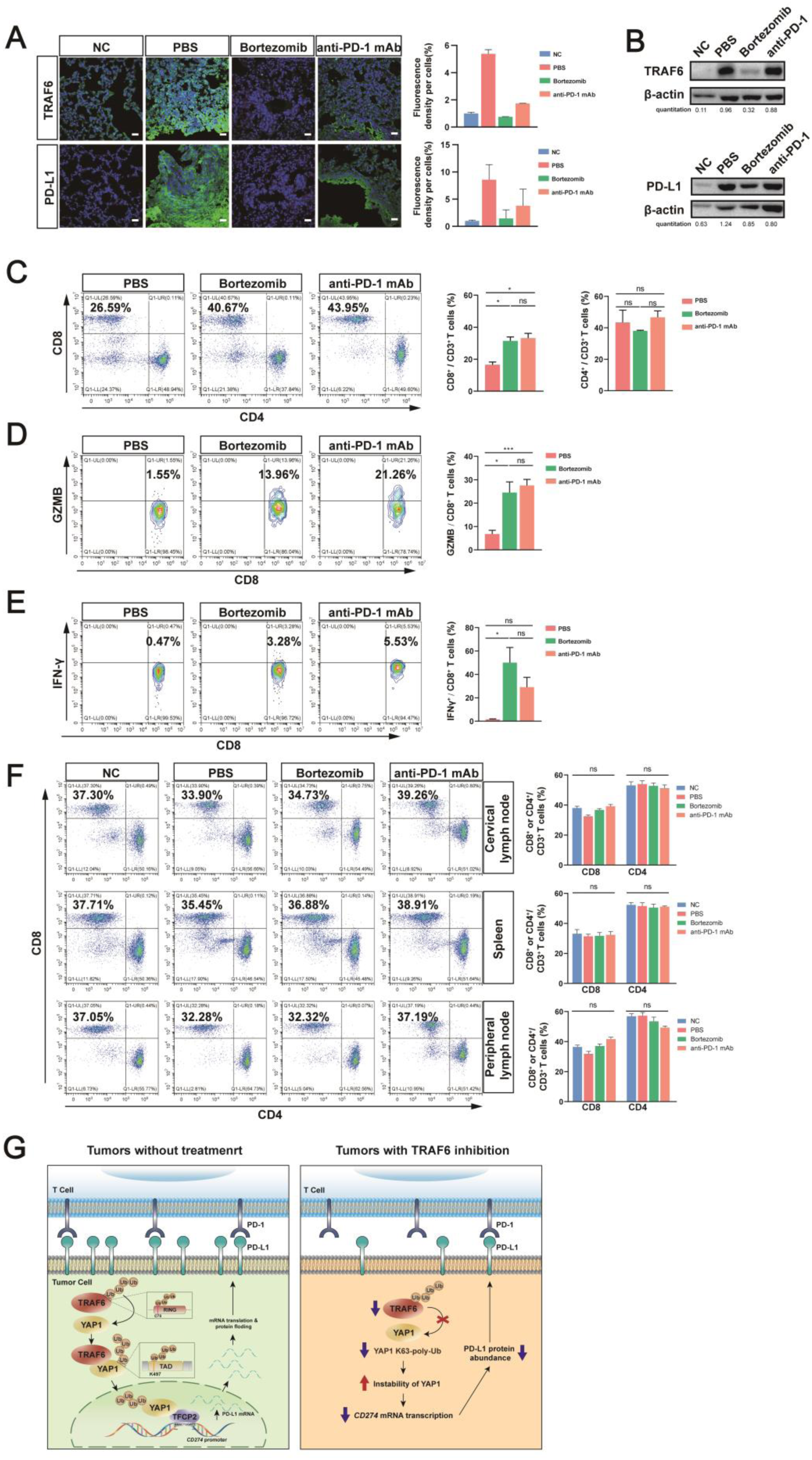
Bortezomib reduced tumor via elevating immune response in situ. **(A)** Representative images of immunofluorescent staining indicated TRAF6 and PD-L1 expression in different therapeutic groups from the lung of mice with B16F10 tumors. Mean with s.d.; n=2. **(B)** Immunoblots analyzed the lung samples carrying B16F10 tumors. **(C-E)** Representative flow cytometric images and quantification of lung samples carrying B16F10 tumors after indicated treatments (C) CD8^+^ T cells and CD4^+^ T cells (D) GZMB^+^ CD8^+^ T cytotoxic cells (E) IFN-γ^+^ CD8^+^ cytotoxic T cells. Mean with s.e.m.; n= 5 mice/group. **(F)** Representative flow cytometric images and quantification of cervical lymph nodes, spleens and peripheral lymph nodes after indicated treatments. Mean with s.e.m.; n=5; In (C-F) *p<0.05; ** p<0.01; ***p<0.001; ns, non-significant; analyzed by Brown-Forsythe and Welch ANOVA tests. **(G)** A working model for targeting TRAF6 illustrated the mechanism that TRAF6 elevated PD-L1 expression via YAP1-TFCP2 composite in melanoma cells. TRAF6 ubiquitinated YAP1 at the K497 site with K63-polyubiquitin chains and further promoted the binding to *CD274 (PD-L1)* promoter of YAP1-TFCP2 composites in the nucleus. This binding facilitates the transcription of *CD274* and increases PD-L1 expression of melanoma cells (left panel). Inhibition of TRAF6 by Bortezomib downregulates PD-L1 and subsequently enhances CD3^+^ cytotoxic T cells driven anti-tumor immunity (right panel).

## Discussion

In this study, we found TRAF6-YAP1-TFCP2 regulating PD-L1, and inhibiting this mechanism reactivate anti-tumor immunity. TRAF6 stabilizes YAP1 by K63 poly-ubiquitination modification, subsequently promoting the formation of YAP1/TFCP2 and PD-L1 transcription. Ubiquitination is an essential type of post-translational modification mediated by the ubiquitin (Ub)-conjugating system composed of E1 Ub-activating enzyme, E2 Ub-conjugating enzyme, and E3 Ub ligase. The K48-linked ubiquitin chain serves as a destruction signal to trigger 26S proteasome-mediated proteolysis, whereas the K63 polyubiquitination plays a role in protein stabilization and function[40]. E3 ligases Cullin 3^SPOP^, β-TrCP and CMTM6 have been reported to destabilize PD-L1 mainly through 26S proteasome- or lysosome-mediated degradation[15, 41–43]. As a special E3 ubiquitin ligase, TRAF6’s RING finger domain can bind to E2 ubiquitin binding enzyme Ubc13/Uev1A to induce K63 polyubiquitination of substrates, stabilize substrate proteins and activate related signaling pathways. Our study revealed that TRAF6 stabilizes YAP1 by catalyzing the K63 polyubiquitylation of YAP1, and the increased YAP1 level can promote PD-L1 transcription after binding with transcription factor TFCP2, thus increasing PD-L1. Although we have confirmed the link of TRAF6 regulating PD-L1, the detailed molecular mechanism remains clarified, including the binding domain between YAP1 and transcription factors in the nucleus, and the specific location of the YAP1-TFCP2 complex binding to PD-L1 promoter region. Understanding these mechanisms provides more possibilities for developing endogenous PD-L1 inhibitors.

Interestingly, TRAF6 is upregulated in various tumors, which has oncogenic characteristics involved in tumorigenesis, tumor development, invasion, and metastasis through various signaling pathways[37–39]. Therefore, targeting TRAF6 has provided a novel strategy for tumor treatment. Small molecular chemicals such as quinine[44], MG132[45] have been reported to interfere with TRAF6, which could down-regulate the expression of TRAF6 in tumors. However, the safety of these chemicals has not been verified clinically. As a first-in-class drug for multiple myeloma, Bortezomib was the first clinically approved proteasome inhibitor, which promote the autophagy mediated lysosomal degradation of TRAF6 and thus down-regulate TRAF6[46]. Therefore, in this study, we chose Bortezomib as the TRAF6 inhibitor to regulate endogenous PD-L1 in melanoma. *In vitro* and *in vivo* experimental results showed that Bortezomib could inhibit tumor TRAF6 expression, down-regulate tumor YAP1 and endogenous PD-L1 expression, and activate tumor-infiltrating CD8^+^ T cells, thus killing melanoma cells more effectively, and had similar efficacy to anti-PD-1 monoclonal antibody therapy. Moreover, 0.2 mg/kg Bortezomib can effectively enhance tumor infiltrating CD8^+^ T cells, and has no significant damage to normal immunity of mice. However, it is important to note that Bortezomib is not a drug that directly interferes with TRAF6 K63 ubiquitination function, suggesting that discovery of more specific inhibitors targeting TRAF6 are necessary. Developing competitive antagonists to block the binding of TRAF6 and E2 ubiquitin binding enzyme, destroying the ubiquitinase activity of TRAF6 without destroying the RING domain, developing protein–protein interaction (PPI) inhibitors to block binding of YAP1 to TRAF6, or the binding between YAP1 and transcription factors may can be used as potential strategies to down-regulate endogenous PD-L1 expression in tumors, providing new ideas and directions for tumor immunotherapy in the future.

## Conclusion

Taken together, we provide direct evidence by a large-scale genetic CRISPR interference (CRISPRi) screen that TRAF6 as a novel driver of PD-L1 in melanoma. Our work also directly reveals that TRAF6 stabilize by catalyzing K63 polyubiquitylation of YAP1, therefore increasing nucleus YAP1 abundance and enhancing the transcriptional activity of YAP1/TFCP2 to induce PD-L1 expression. Of importance, TRAF6 has also shown the potential as a sensitive biomarker to evaluate drug responses of PD-1 inhibitors in the preclinical metastatic melanoma model. Hence, our study not only provides a molecular insight, but also reveals a potential therapeutic strategy that targeting the E3 ligase TRAF6 as an endogenous down-regulation strategy might enhance the efficacy of anti-PD-1/PD-L1 in treating human cancers.

## Availability of data and materials

The datasets sharing used and/or analyzed during the current study are available from the corresponding author Fang Cheng on reasonable request.

## Abbreviation

CRISPR: Clustered Regularly Interspaced Short Palindromic Repeats
TRAF6: Tumor Necrosis Factor Receptor Associated Factor 6
PD-L1: Programmed death ligand 1
PD-L2: Programmed death ligand 2
YAP1: Yes-associated protein 1
TFCP2: Transcription Factor CP2
JAK2: Janus Kinase 2
PTEN: Phosphatase and Tensin Homolog
PI3K: Phosphoinositide 3-kinase
EGFR: Epidermal Growth Factor Receptor
CDK5: Cyclin Dependent Kinase 5
IFN-γ: IFN-gamma
mAb: monoclonal antibody

## Author Contribution

Xiaoyan Liu and Linglu Wang contributed equally to this manuscript.

## Contributions

FC, XYL and LLW conceptualized and designed the research work. HTZ and HBC provided conceptual advice and supervised the study. XYL, LLW, YHH and FS performed the experiments, data curation and original draft preparation. HT and HC contributed to the design of experiments. ZXX contributed to the CRIPSRi library screen work. HZ, HBC, FC performed project administration and funding acquisition. FC finalized the manuscript with the contributions of other authors. All authors read and approved the final manuscript.

## Corresponding authors

**Correspondence to** Haitao Zhu, Hongbo Chen, Fang Cheng

## Ethics declarations

### Ethics approval and consent to participate

The use of laboratory animals and all animal experiments was reviewed and approved by the Institutional Animal Care and Use Committee (IACUC), Sun Yat-Sen University, China. The approval number is SYSU-IACUC-2020-000474.

### Consent for publication

Not applicable.

## Supplementary Information

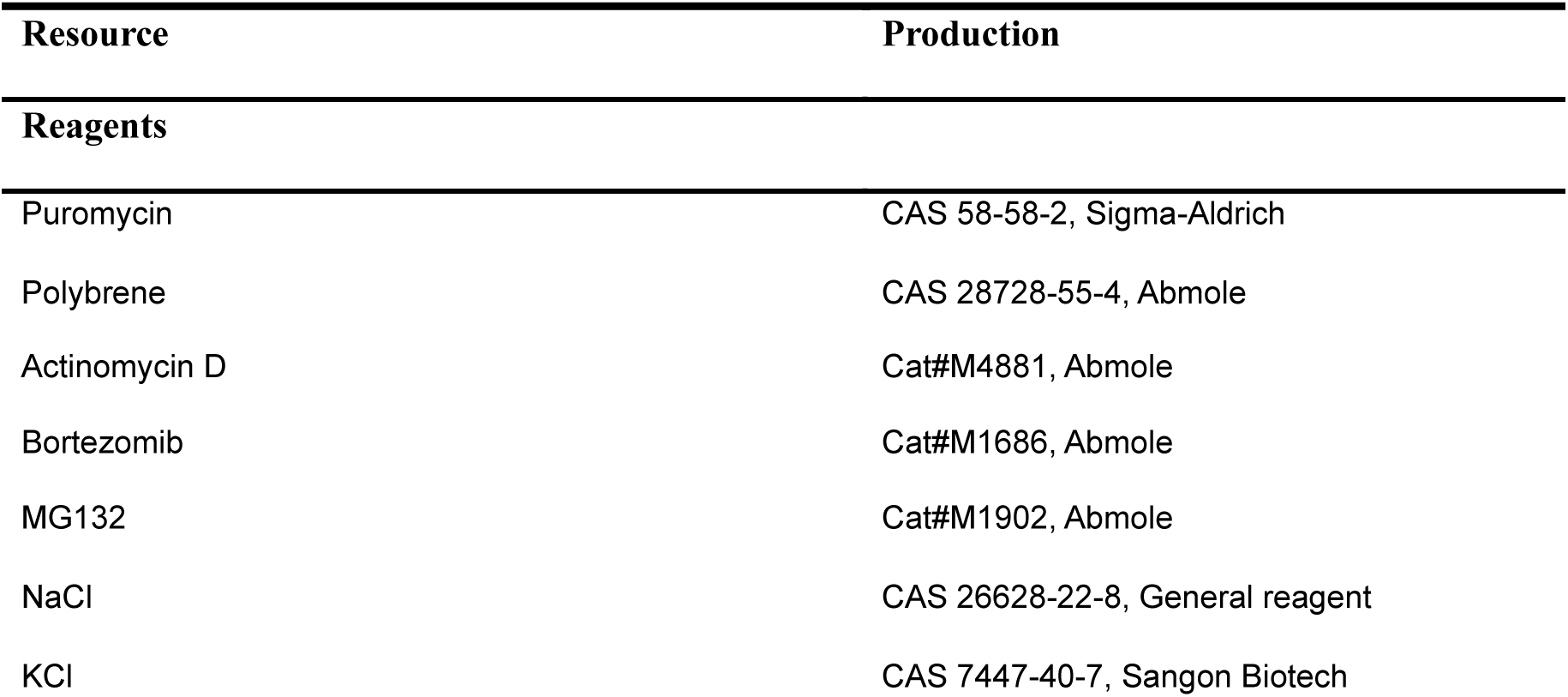
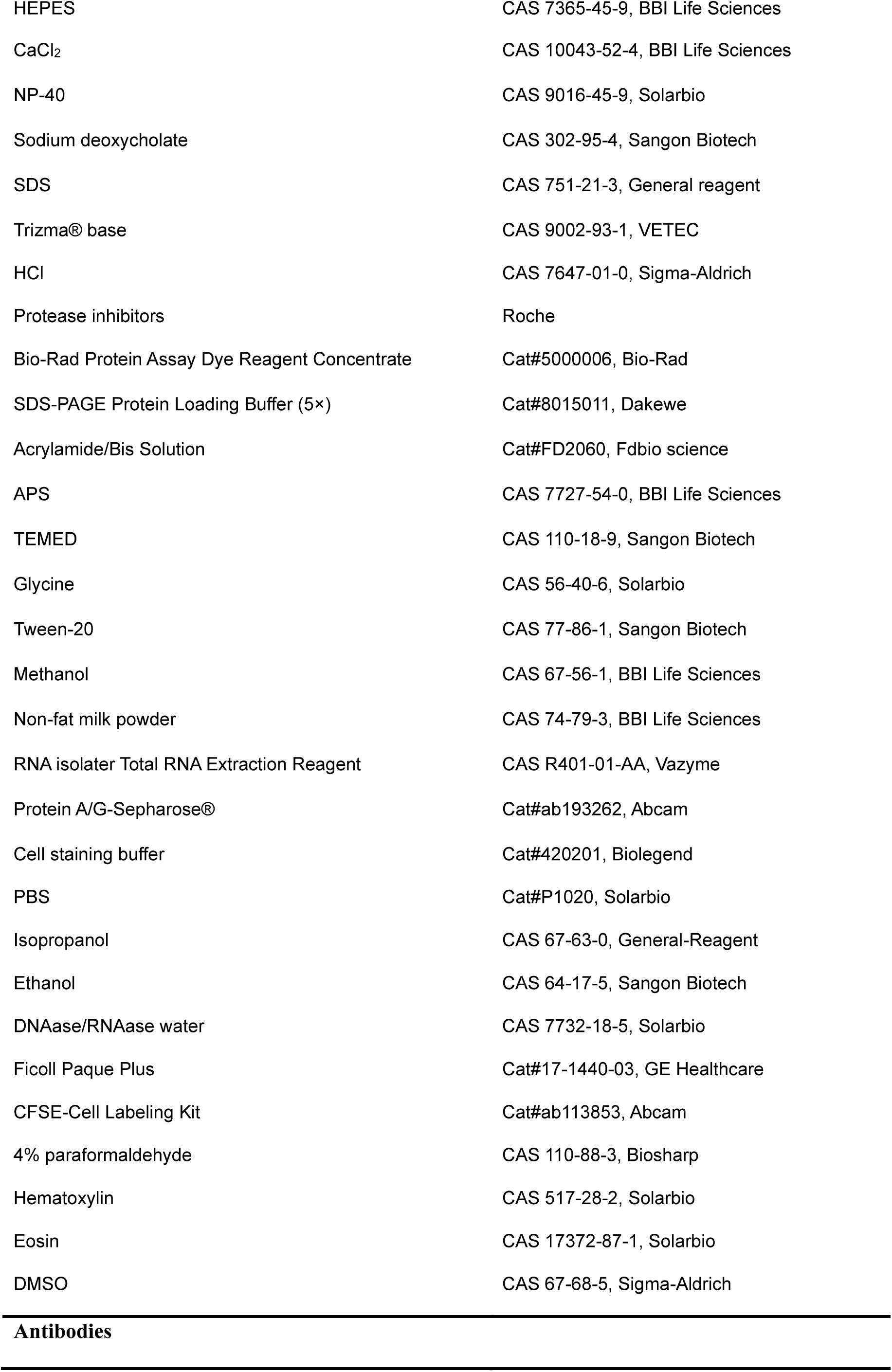
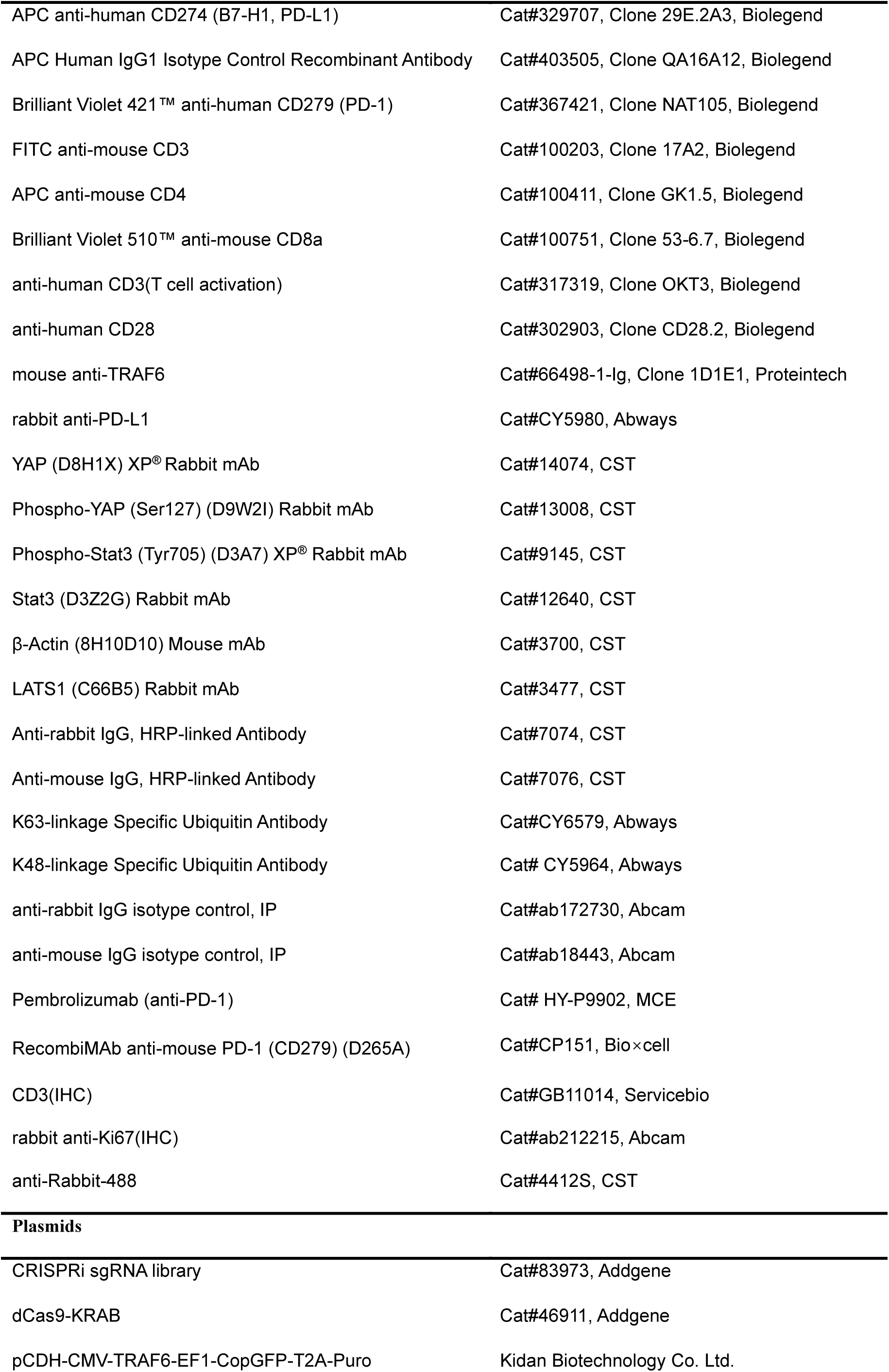
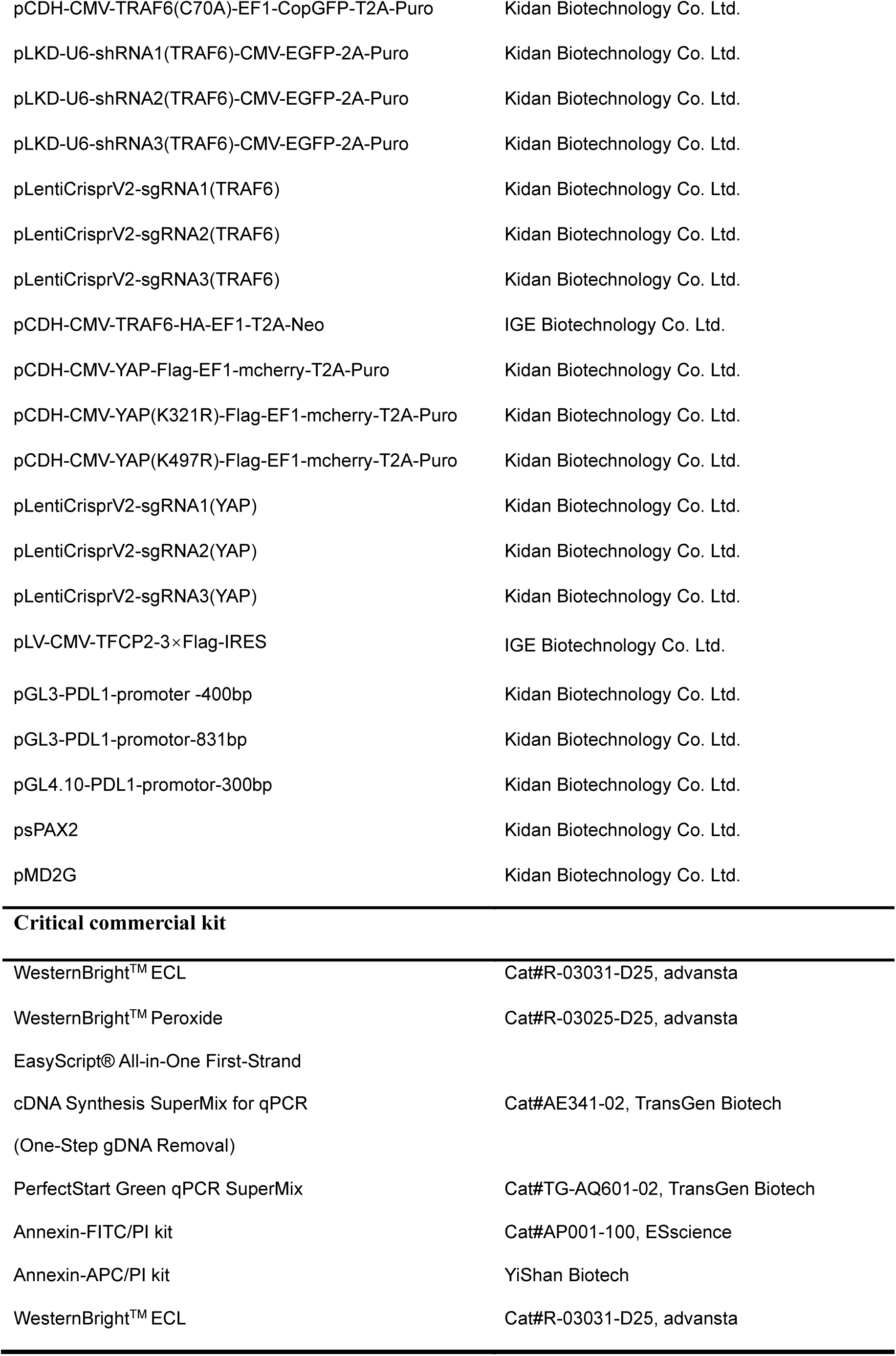
Supplementary Table 1.

Supplementary Table 1.
| Resource | Production |
| --- | --- |
| Reagents |  |
| Puromycin | CAS 58-58-2, Sigma-Aldrich |
| Polybrene | CAS 28728-55-4, Abmole |
| Actinomycin D | Cat#M4881, Abmole |
| Bortezomib | Cat#M1686, Abmole |
| MG132 | Cat#M1902, Abmole |
| NaCl | CAS 26628-22-8, General reagent |
| KCl | CAS 7447-40-7, Sangon Biotech |
| HEPES | CAS 7365-45-9, BBI Life Sciences |
| CaCl<sub>2</sub> | CAS 10043-52-4, BBI Life Sciences |
| NP-40 | CAS 9016-45-9, Solarbio |
| Sodium deoxycholate | CAS 302-95-4, Sangon Biotech |
| SDS | CAS 751-21-3, General reagent |
| Trizma® base | CAS 9002-93-1, VETEC |
| HCl | CAS 7647-01-0, Sigma-Aldrich |
| Protease inhibitors | Roche |
| Bio-Rad Protein Assay Dye Reagent Concentrate | Cat#5000006, Bio-Rad |
| SDS-PAGE Protein Loading Buffer (5×) | Cat#8015011, Dakewe |
| Acrylamide/Bis Solution | Cat#FD2060, Fdbio science |
| APS | CAS 7727-54-0, BBI Life Sciences |
| TEMED | CAS 110-18-9, Sangon Biotech |
| Glycine | CAS 56-40-6, Solarbio |
| Tween-20 | CAS 77-86-1, Sangon Biotech |
| Methanol | CAS 67-56-1, BBI Life Sciences |
| Non-fat milk powder | CAS 74-79-3, BBI Life Sciences |
| RNA isolater Total RNA Extraction Reagent | CAS R401-01-AA, Vazyme |
| Protein A/G-Sepharose® | Cat#ab193262, Abcam |
| Cell staining buffer | Cat#420201, Biolegend |
| PBS | Cat#P1020, Solarbio |
| Isopropanol | CAS 67-63-0, General-Reagent |
| Ethanol | CAS 64-17-5, Sangon Biotech |
| DNAase/RNAase water | CAS 7732-18-5, Solarbio |
| Ficoll Paque Plus | Cat#17-1440-03, GE Healthcare |
| CFSE-Cell Labeling Kit | Cat#ab113853, Abcam |
| 4% paraformaldehyde | CAS 110-88-3, Biosharp |
| Hematoxylin | CAS 517-28-2, Solarbio |
| Eosin | CAS 17372-87-1, Solarbio |
| DMSO | CAS 67-68-5, Sigma-Aldrich |
| Antibodies |  |
| APC anti-human CD274 (B7-H1, PD-L1) | Cat#329707, Clone 29E.2A3, Biolegend |
| APC Human IgG1 Isotype Control Recombinant Antibody | Cat#403505, Clone QA16A12, Biolegend |
| Brilliant Violet 421™ anti-human CD279 (PD-1) | Cat#367421, Clone NAT105, Biolegend |
| FITC anti-mouse CD3 | Cat#100203, Clone 17A2, Biolegend |
| APC anti-mouse CD4 | Cat#100411, Clone GK1.5, Biolegend |
| Brilliant Violet 510™ anti-mouse CD8a | Cat#100751, Clone 53-6.7, Biolegend |
| anti-human CD3(T cell activation) | Cat#317319, Clone OKT3, Biolegend |
| anti-human CD28 | Cat#302903, Clone CD28.2, Biolegend |
| mouse anti-TRAF6 | Cat#66498-1-Ig, Clone 1D1E1, Proteintech |
| rabbit anti-PD-L1 | Cat#CY5980, Abways |
| YAP (D8H1X) XP® Rabbit mAb | Cat#14074, CST |
| Phospho-YAP (Ser127) (D9W2I) Rabbit mAb | Cat#13008, CST |
| Phospho-Stat3 (Tyr705) (D3A7) XP® Rabbit mAb | Cat#9145, CST |
| Stat3 (D3Z2G) Rabbit mAb | Cat#12640, CST |
| β-Actin (8H10D10) Mouse mAb | Cat#3700, CST |
| LATS1 (C66B5) Rabbit mAb | Cat#3477, CST |
| Anti-rabbit IgG, HRP-linked Antibody | Cat#7074, CST |
| Anti-mouse IgG, HRP-linked Antibody | Cat#7076, CST |
| K63-linkage Specific Ubiquitin Antibody | Cat#CY6579, Abways |
| K48-linkage Specific Ubiquitin Antibody | Cat# CY5964, Abways |
| anti-rabbit IgG isotype control, IP | Cat#ab172730, Abcam |
| anti-mouse IgG isotype control, IP | Cat#ab18443, Abcam |
| Pembrolizumab (anti-PD-1) | Cat# HY-P9902, MCE |
| RecombiMAb anti-mouse PD-1 (CD279) (D265A) | Cat#CP151, Bio×cell |
| CD3(IHC) | Cat#GB11014, Servicebio |
| rabbit anti-Ki67(IHC) | Cat#ab212215, Abcam |
| anti-Rabbit-488 | Cat#4412S, CST |
| **Plasmids** |  |
| CRISPRi sgRNA library | Cat#83973, Addgene |
| dCas9-KRAB | Cat#46911, Addgene |
| pCDH-CMV-TRAF6-EF1-CopGFP-T2A-Puro | Kidan Biotechnology Co. Ltd. |
| pCDH-CMV-TRAF6(C70A)-EF1-CopGFP-T2A-Puro | Kidan Biotechnology Co. Ltd. |
| pLKD-U6-shRNA1(TRAF6)-CMV-EGFP-2A-Puro | Kidan Biotechnology Co. Ltd. |
| pLKD-U6-shRNA2(TRAF6)-CMV-EGFP-2A-Puro | Kidan Biotechnology Co. Ltd. |
| pLKD-U6-shRNA3(TRAF6)-CMV-EGFP-2A-Puro | Kidan Biotechnology Co. Ltd. |
| pLentiCrisprV2-sgRNA1(TRAF6) | Kidan Biotechnology Co. Ltd. |
| pLentiCrisprV2-sgRNA2(TRAF6) | Kidan Biotechnology Co. Ltd. |
| pLentiCrisprV2-sgRNA3(TRAF6) | Kidan Biotechnology Co. Ltd. |
| pCDH-CMV-TRAF6-HA-EF1-T2A-Neo | IGE Biotechnology Co. Ltd. |
| pCDH-CMV-YAP-Flag-EF1-mcherry-T2A-Puro | Kidan Biotechnology Co. Ltd. |
| pCDH-CMV-YAP(K321R)-Flag-EF1-mcherry-T2A-Puro | Kidan Biotechnology Co. Ltd. |
| pCDH-CMV-YAP(K497R)-Flag-EF1-mcherry-T2A-Puro | Kidan Biotechnology Co. Ltd. |
| pLentiCrisprV2-sgRNA1(YAP) | Kidan Biotechnology Co. Ltd. |
| pLentiCrisprV2-sgRNA2(YAP) | Kidan Biotechnology Co. Ltd. |
| pLentiCrisprV2-sgRNA3(YAP) | Kidan Biotechnology Co. Ltd. |
| pLV-CMV-TFCP2-3×Flag-IRES | IGE Biotechnology Co. Ltd. |
| pGL3-PDL1-promoter -400bp | Kidan Biotechnology Co. Ltd. |
| pGL3-PDL1-promotor-831bp | Kidan Biotechnology Co. Ltd. |
| pGL4.10-PDL1-promotor-300bp | Kidan Biotechnology Co. Ltd. |
| psPAX2 | Kidan Biotechnology Co. Ltd. |
| pMD2G | Kidan Biotechnology Co. Ltd. |
| Critical commercial kit |  |
| WesternBright™ ECL | Cat#R-03031-D25, advansta |
| WesternBright™ Peroxide | Cat#R-03025-D25, advansta |
| EasyScript® All-in-One First-Strand |  |
| cDNA Synthesis SuperMix for qPCR<br>(One-Step gDNA Removal) | Cat#AE341-02, TransGen Biotech |
| PerfectStart Green qPCR SuperMix | Cat#TG-AQ601-02, TransGen Biotech |
| Annexin-FITC/PI kit | Cat#AP001-100, ESscience |
| Annexin-APC/PI kit | YiShan Biotech |
| WesternBright™ ECL | Cat#R-03031-D25, advansta |

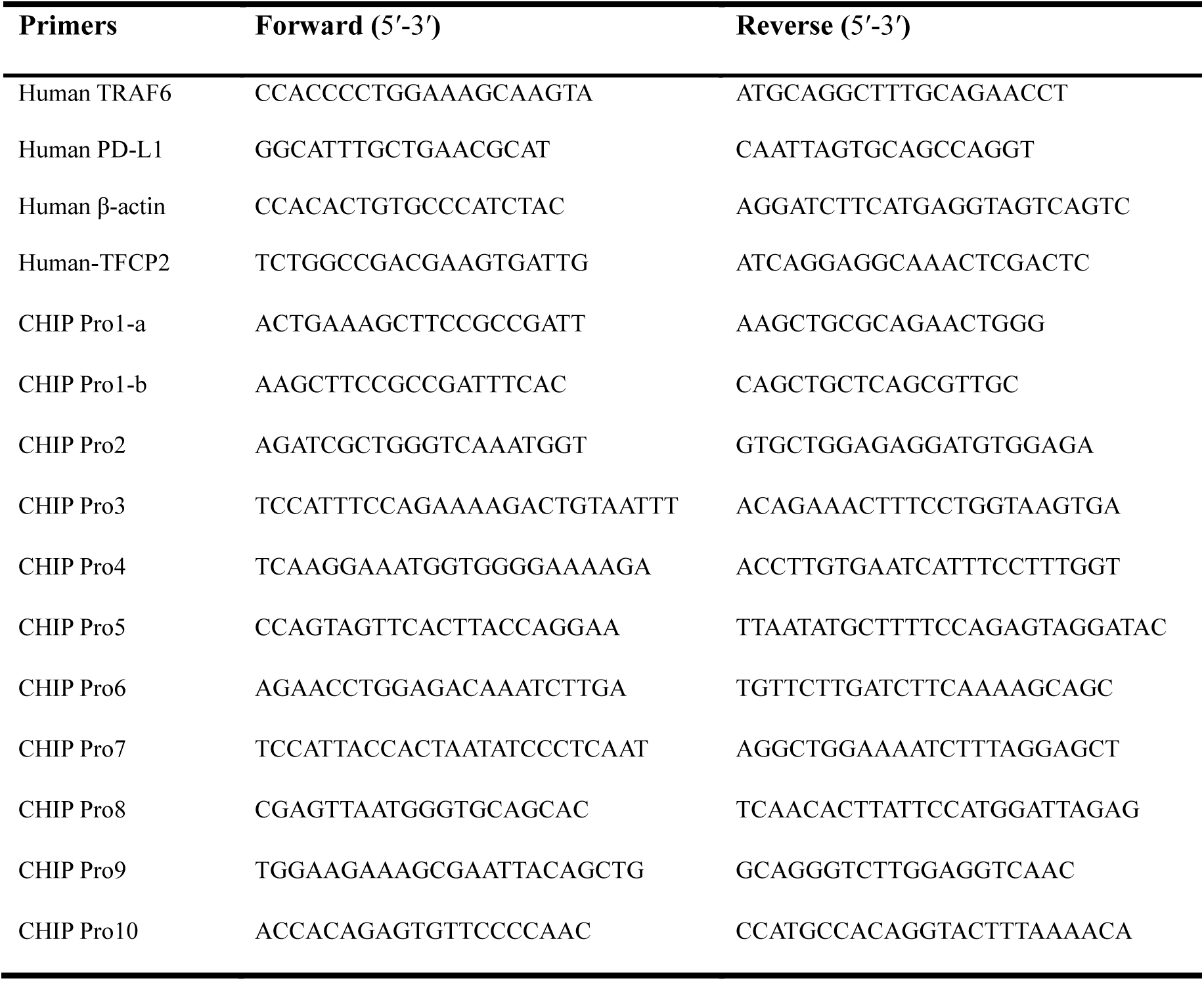
Supplementary Table 2

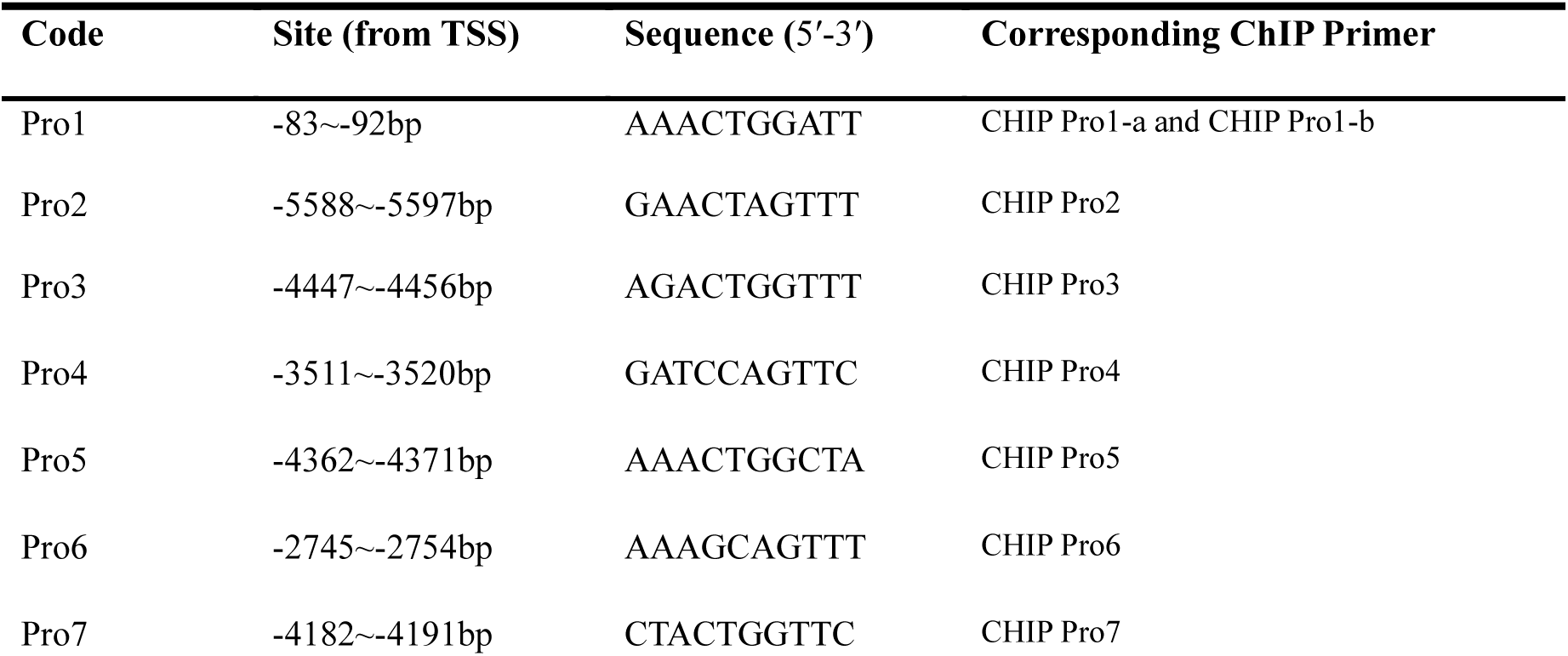
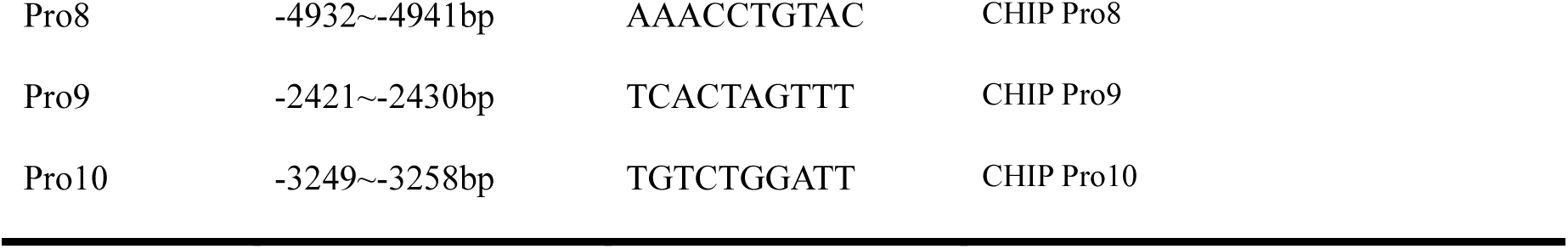
Supplementary Table 3

